# Troriluzole Rescues Glutamatergic Deficits, Amyloid and Tau Pathology, and Synaptic and Memory Impairments in 3xTg-AD Mice

**DOI:** 10.1101/2024.06.11.598453

**Authors:** Jeremiah Pfitzer, Priyanka D. Pinky, Savannah Perman, Emma Redmon, Luca Cmelak, Vishnu Suppiramaniam, Vladimir Coric, Irfan A. Qureshi, Michael W. Gramlich, Miranda N. Reed

**Affiliations:** Department of Drug Discovery and Development, Auburn University, Auburn, AL, USA; Department of Psychological Sciences, Auburn University, Auburn, AL, USA; Center for Neuroscience Initiative, Auburn University, Auburn, AL, USA; Department of Molecular and Cellular Biology, College of Science and Mathematics, Kennesaw State University, Kennesaw, Georgia; Biohaven Pharmaceuticals Inc., New Haven, CT, USA; Department of Physics, Auburn University, Auburn, AL, USA

**Keywords:** Alzheimer’s disease, glutamate, amyloid, tau, vesicular glutamate transporter, presynaptic, synaptic plasticity, memory

## Abstract

Alzheimer’s disease (AD) is a neurodegenerative condition in which clinical symptoms are highly correlated with the loss of glutamatergic synapses. While later stages of AD are associated with markedly decreased glutamate levels due to neuronal loss, in the early stages, pathological accumulation of glutamate and hyperactivity contribute to AD pathology and cognitive dysfunction. There is increasing awareness that presynaptic dysfunction, particularly synaptic vesicle (SV) alterations, play a key role in mediating this early-stage hyperactivity. In the current study, we sought to determine whether the 3xTg mouse model of AD that exhibits both beta-amyloid (Aβ) and tau-related pathology would exhibit similar presynaptic changes as previously observed in amyloid or tau models separately. Hippocampal cultures from 3xTg mice were used to determine whether presynaptic vesicular glutamate transporters (VGlut) and glutamate are increased at the synaptic level while controlling for postsynaptic activity. We observed that 3xTg hippocampal cultures exhibited increased VGlut1 associated with an increase in glutamate release, similar to prior observations in cultures from tau mouse models. However, the SV pool size was also increased in 3xTg cultures, an effect not previously observed in tau mouse models but observed in Aβ models, suggesting the changes in pool size may be due to Aβ and not tau. Second, we sought to determine whether treatment with troriluzole, a novel 3^rd^ generation tripeptide prodrug of the glutamate modulator riluzole, could reduce VGlut1 and glutamate release to restore cognitive deficits in 8-month-old 3xTg mice. Treatment with troriluzole reduced VGlut1 expression, decreased basal and evoked glutamate release, and restored cognitive deficits in 3xTg mice. Together, these findings suggest presynaptic alterations are early events in AD that represent potential targets for therapeutic intervention, and these results support the promise of glutamate-modulating drugs such as troriluzole in Alzheimer’s disease.

## Introduction

Alzheimer’s disease (AD) is a neurodegenerative condition and the most common cause of dementia. The hallmarks of AD include extracellular deposition of β-amyloid (Aβ) in the form of diffuse and neuritic plaques and the presence of intraneuronal neurofibrillary tangles consisting of aggregated hyperphosphorylated tau protein, as well as neuronal loss (Duyckaerts *et al*. 2009). Before neuronal level loss, however, synaptic level dysfunction and loss are often observed in the prodromal and early stages of AD (Pelucchi *et al*. 2022). Notably, clinical symptoms in AD patients are highly correlated with the loss of glutamatergic synapses (Overk *et al*. 2014; Terry *et al*. 1991), which play a critical role in mediating memory. Glutamate is the major excitatory neurotransmitter in the brain, and over 40% of neuronal synapses are glutamatergic. In some regions, such as the cerebral cortex and hippocampus, 80-90% of neurons use glutamate as a neurotransmitter (Somogyi *et al*. 1998), with the hippocampus being a critical brain region for learning and memory and one of the first to be affected in AD. While later stages of AD are associated with markedly decreased glutamate levels due to neuronal loss, in the early stages, pathological accumulation of glutamate and hyperactivity contribute to AD pathology and cognitive dysfunction (Cox *et al*. 2022; Pinky *et al*. 2023b; Rudy *et al*. 2014; Setti & Reed 2022). While much of this research has examined disruptions in postsynaptic responses, there is increasing awareness of the role that presynaptic dysfunction plays in mediating hyperactivity (Wu *et al*. 2021). For example, a recent study with two mouse models of preclinical AD revealed selective impairment of hippocampal synaptic vesicle (SV) proteins before detectable increases in Aβ_42_ levels (Hark *et al*. 2021). This impaired turnover of SVs resulted in elevated levels, enlargement of the SV pool, and enhancement of presynaptic potentiation. Presynapses being an early site of dysfunction is not necessarily surprising considering that the localization and processing of amyloid precursor protein (APP) mainly occur at presynaptic terminals, and APP is trafficked and secreted via SVs (Hark *et al*. 2021; Groemer *et al*. 2011). These changes may be permissive for the spread of pathological Aβ and tau, as glutamate release or its synaptic activity have been shown to induce the release of both into the extracellular space (Bero *et al*. 2011; de Calignon *et al*. 2012; Pooler *et al*. 2013; Yamada *et al*. 2014; Kamenetz *et al*. 2003; Dani *et al*. 2016).

Aβ has been shown to preferentially accumulate in presynaptic terminals that contain the vesicular glutamate transporter VGlut1 in AD patients (Sokolow *et al*. 2012), suggesting Aβ may preferentially affect presynaptic glutamatergic terminals. Though VGlut1 is decreased in the later stages of AD (Kashani *et al*. 2008), patients with mild cognitive impairment (MCI) exhibit increased VGlut1 (Bell *et al*. 2007). This increase in VGlut1 may help explain why hippocampal hyperexcitability is observed in MCI patients, as increasing number of vesicular transporters results in enhanced evoked and miniature responses, as well as an increase in the amount of glutamate released per vesicle into the synaptic cleft (Wilson *et al*. 2005). Interestingly, the degree of hyperexcitability in MCI patients predicts the conversion to AD and the rate of cognitive decline (Mackenzie & Miller 1994). The APP/PS1 mouse model of AD also exhibits an early increase in VGlut1 expression concurrent with elevated extracellular glutamate levels, as well as stimulus-evoked glutamate release, that precedes cognitive deficits (Hascup *et al*. 2019; Hascup & Hascup 2015). Treatment with riluzole, a protein kinase C inhibitor that reduces glutamate release, resulted in decreased basal glutamate and stimulus-evoked glutamate release and improved cognitive functioning despite no change in Aβ plaque accumulation (Hascup *et al*. 2021).

The PS19 tauopathy mouse model also exhibits an early increase followed by a later decrease in VGlut1 expression pattern (Crescenzi *et al*. 2017). We have previously shown *in vivo* that memory deficits in the rTg(TauP301L)4510 mouse model were associated with a greater than 40% increase in VGlut1 levels and a corresponding increase in glutamate release in the hippocampus at an age when there is no detectable neuron loss and only subtle tau pathology (Hunsberger *et al*. 2014; Hunsberger *et al*. 2021). Reducing VGlut1 levels and glutamate release with riluzole treatment restored memory deficits in the rTg4510 model (Hunsberger *et al*. 2015c). However, still unclear was whether VGlut1 levels mediated increased extracellular glutamate, as changes in VGlut1 levels have been shown to directly affect the presynaptic release probability (Wilson *et al*. 2005). To address this question, we used hippocampal cultures from rTg4510 mice and a pH-sensitive green fluorescent protein approach; cultures from rTg4510 mice exhibited a 40% increase in VGlut1 levels, as well as a 40% increase in glutamate release, that was not caused by increased vesicle exocytosis probability nor an increase in vesicle pool size but rather was directly related to increased VGlut1 transporters per synaptic vesicle (Taipala *et al*. 2022). This is notable as APP knock-in mice exhibit an enlargement in the SV pool during the early stages of Aβ proteotoxicity before synaptic transmission becomes impaired (Hark *et al*. 2021), suggesting Aβ and tau may have differential effects on presynaptic terminals.

The purpose of the current studies was two-fold. First, we sought to determine whether the 3xTg mouse model of AD that has three mutations associated with familial Alzheimer’s disease (APP Swedish, MAPT P301L, and PSEN1 M146V) and thus exhibit both Aβ and tau-related pathology would exhibit similar changes in VGlut1 and glutamate release as previously observed in APP/PS1 and rTg4510 mice separately. To achieve this goal, we examined hippocampal cultures from these mice to determine whether presynaptic VGlut1 and glutamate levels are increased at the synaptic level while controlling for postsynaptic activity. We then imaged VGlut1 expression per SV, presynaptic SV release mechanics, presynaptic recycling pool size, and total presynaptic SV pool size. We were then able to quantify all metrics in the same presynapses during the same experiment. Second, we sought to determine whether treatment with troriluzole, a novel 3^rd^ generation tripeptide prodrug of the glutamate modulator riluzole, could reduce VGlut1 and glutamate release to restore cognitive deficits in 8-month-old 3xTg mice. As a rationally designed prodrug of riluzole, troriluzole was engineered to overcome some of the pharmaceutical limitations of riluzole, including its low bioavailability, variable pharmacokinetic profile, negative food effect, requirement for twice daily dosing, and potential for hepatotoxicity (Pelletier *et al*. 2018; Pelletier *et al*. 2014; McDonnell *et al*. 2012). We observed that 3xTg hippocampal cultures did exhibit increased VGlut1 associated with an increase in glutamate release, similar to what we previously observed in rTg4510 cultures (Taipala *et al*. 2022). Interestingly, the SV pool size was also increased in 3xTg cultures, an effect not previously observed in the rTg4510 model (Taipala *et al*. 2022) but observed in APP knock-in mice (Hark *et al*. 2021), suggesting the changes in pool size may be due to Aβ and not tau. We also observed that treatment with troriluzole reduced VGlut1 expression, decreased basal and evoked glutamate release, and restored cognitive deficits in 3xTg mice. Together, these findings suggest presynaptic alterations are early events in AD that represent potential for therapeutic intervention.

## Material & Methods

### Mice

Female triple transgenic Alzheimer’s disease mice (3xTg-AD, MMRRC stock #34830) expressing amyloid precursor protein with the Swedish familial double mutation (APP_KM670/671NL_), presenilin 1 with a substitution mutation (PSEN1_M146V_), and microtubule-associated protein tau mutation (MAPT_P301L_) were initially purchased from Jackson’s lab and bred in-house, as previously described (Oddo *et al*. 2003). Experimental 3xTg-AD mice were positively genotyped at the beginning and end of the study for amyloid precursor protein Swedish (APPswe) KM670/671NL, presenilin 1 (PSEN1) M146V, and microtubule-associated protein tau (MAPT) P301L mutations (Sterniczuk *et al*. 2010). Only female mice were used in the current study due to the lack of a robust phenotype in male mice (Clinton *et al*. 2007). Nontransgenic mice (TgNeg; stock #101043) of the same background strain (129/C57BL6) were also purchased from The Jackson Laboratory and bred in-house. All mice were group housed, provided food and water *ad libitum,* and housed under 12:12 hour light-dark cycles in humidity and temperature-controlled rooms. All experiments were performed during the light cycle. The Auburn University Animal Care and Use Committee approved all experiments (protocol numbers: 2018-3408 & 2020-3716), which complied with the guidelines published in the NIH Guide for the Care and Use of Laboratory Animals.

### Cell Cultures

All animal dissections occurred on postnatal day 4 or 5 pups that were genotyped before dissection to confirm gene expression. Cell cultures were plated on glass coverslips in 12-well plates according to established methods (Peng *et al*. 2012). First, a layer of primary astrocytes was plated and allowed to achieve uniform confluence for one week in fetal calf serum. Then, hippocampi from pups were dissected, combined, and plated in neuronal media. Because of the early age, sex could not be determined, and thus, pups of equal sexes were assumed. All neurons were plated at a density of 20k per plate, consistent with our previously established density for single presynaptic VGlut1 imaging (Taipala *et al*. 2022). Twelve hours after plating, cell media was changed to a fresh neurobasal medium for neuronal growth. Cultures were then incubated for 14-21 Days In Vitro (DIV) with media replacement every five days.

### VGlut1-pHluorin Transfection

VGlut1-pHluorin was generously provided by Drs. Robert Edward and Susan Voglmaier (UCSF) (Voglmaier et al., 2006). Lentiviral vectors were generated by the Viral Vectors Core at Washington University and utilized in accordance with approved procedures at Auburn University. Cultures were transfected at DIV3 for 48 hrs. The media was then replaced with fresh neurobasal media, and half media replacements occurred every three days until the day of imaging. During imaging, samples were exposed to imaging media containing 140 mM NaCl, 2.5 mM KCl, 2 mM CaCl2, 4 mM MgCl2, 10 mM HEPES, 2 mM Glucose, 50 mM DL-AP5, 10 mM CNQX, pH adjusted to pH 7.4, following our previously established protocol.

### VGlut1-pHluorin Imaging and Data Analysis

Cell culture measurements were performed with a Nikon Ti-2 microscope (Nikon) using an EPI fluorescent GFP cube (Nikon) with a 100x Oil immersion objective (Nikon) and ORCA Flash v4 CMOS camera (Hamamatsu), with a pixel resolution of 0.065 x 0.065 mm. The culture temperature was held at 37 °C during experiments. Cultures were stimulated with a uniform electrical field using two parallel electrodes. VGlut1-pHluorin stimulation protocol was performed as described in the results section. Individual frames from VGlut1-pHluorin movies were background subtracted using a 10-frame rolling-ball-radius package in ImageJ. Presynapses were then identified during 40 Hz stimulation using in-lab Matlab scripts. Identified presynapse location intensities were then run through a Kennedy-Chung filter algorithm previously shown to identify spontaneous release events (Chanaday & Kavalali, 2018). We then counted intensity spikes following our previously established approach of at least one standard deviation above the noise level (Taipala et al., 2022). The number of identified release events was then aggregated and divided by the observation time to estimate release frequency. The spontaneous and stimulate release distributions were then fit to Poisson distribution 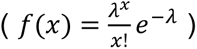 to determine the mean value (λ). The intensity per vesicle was determined as the three-frame average of the peak minus the three-frame average of the baseline before release.

### Computational Model of Presynaptic Release

Computational simulations were based on our previously established binomial model (Taipala et al., 2022) and previous presynaptic release models (Lanore & Silver, 2016; Reid & Clements, 1999; Scheuss & Neher, 2001). Here, we performed a multinomial vesicle release model of multiple release sites described as: *I*(*t_i_*) = *Npq*Θ(*t* − *t*_0_) + *I*(*t*_(*i*−1)_) where N is the number of active zone sites, p is the probability of release per site (assumed to be constant for all sites), q is the intensity per vesicle. All model parameters used to reproduce each experimental condition were constrained. We also used a coarse-grained exponentially decaying vesicle release probability 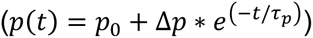 to represent combined presynaptic factors that have been extensively explored but are beyond the scope of this present study. Simulations were performed using Python 3.10 release, and the algorithm follows our previous approach (Taipala et al., 2022).

### Troriluzole administration

For *in vivo* experiments, TgNeg and 3xTg mice were assigned at five months of age via simple randomization to receive either 25 mg/kg/day troriluzole (Tro; Biohaven Pharmaceuticals) + 1% w/v saccharin (vehicle) dissolved in drinking water or vehicle (Veh) alone. There were three groups: vehicle-treated transgene-negative littermates (Veh-TgNeg, n = 19), vehicle-treated 3xTg mice (Veh-3xTg, n = 14), and troriluzole-treated 3xTg mice (Tro-3xTg, n = 15). A sample size of 14 per group was derived from a power analysis (power = 0.8, *p*<0.05; two-tailed; expected Cohen’s d > 0.8 or f >0.4) from preliminary behavioral studies comparing TgNegs and 3xTg. Troriluzole was dissolved by stirring the compound in room-temperature DI water. Water consumption was measured, and concentrations were adjusted every 3-5 days to ensure that intake remained at ∼25 mg/kg/day per mouse, approximately equivalent to riluzole used in previous studies (Okamoto *et al*. 2018; Ishiyama *et al*. 2004; Gourley *et al*. 2012; Hunsberger *et al*. 2015c; Hunsberger *et al*. 2016a). To account for water loss during routine cage handling, a control cage was handled the same as the treatment cages, and this cage’s water loss was measured and subtracted from dosing calculations. Administration via drinking water was implemented to reduce the stress associated with daily handling that occurs with other methods, such as daily gavage, and to allow for comparison to prior riluzole studies using this route of administration in preclinical models (Hunsberger *et al*. 2016a; Hunsberger *et al*. 2015c; Okamoto *et al*. 2018; Schmidt *et al*. 2016; Lesuis *et al*. 2019).

### Morris water maze

At eight months of age, after three months of treatment, mice underwent training in the Morris water maze using our established protocol (Weitzner *et al*. 2015). A circular pool 100 centimeters in diameter was filled with water maintained at 20 ± 1°C. Mice were subjected to 8 days of Morris water maze (MWM) testing, including four preliminary visual platform training trials on day 1, four hidden platform training trials on days 2-7, and probe trials on the start of days 4-6 and 8, as previously described (Weitzner *et al*. 2015). Swim speed and path length were quantified for visible and hidden platform training. The percentage of time spent in the target quadrant was quantified for probe trials using EthoVision XT (Noldus, VA) software.

On the first day, mice underwent visual platform training to ensure any deficits observed in the hidden platform or probe trials of the MWM were not due to visual or motor deficits. Mice were placed in the pool facing opposite the rescue platform. Mice were given 60 seconds to swim to a visible platform that was elevated 1 centimeter above the water and further indicated with a flag. If the mice did not find the platform after 60 seconds, they were guided to it. The mice remained on the platform for 15 seconds before being removed, dried, and placed in a holding cage with access to a heating pad. The mice underwent four trials with an inter-trial interval of approximately 15 minutes. The platform location was moved between each trial.

On days 2-7, mice were trained to swim to a hidden platform 9 centimeters in diameter located 1 centimeter below the water in one of the four quadrants. The pool was made opaque with non-toxic white paint, and visual cues were located around the room. The mice were placed in the pool at predetermined, pseudorandom locations and swam until they reached the platform or 60 seconds passed. If mice did not find the platform, they were gently guided to it. After reaching the platform, all mice remained for 15 seconds and were dried off and placed in a cage with access to a heating pad. The mice were trained four times daily on days 2-7 with an inter-trial interval of 20 minutes.

Prior to the first hidden platform trials on days 4, 5, 6, and 8 of MWM testing, mice underwent a 60-second probe trial in which the platform was removed, and mice swam freely in the pool. The percentage of time in the target quadrant was defined as the time spent in the quadrant where the rescue platform was located divided by the total time and was compared. A percentage time of 25% in the target quadrant indicates chance performance.

### Elevated plus maze

Twenty-four hours following the completion of the Morris water maze, mice were placed in the center of a plus-shaped maze with four arms. Two of the maze’s opposing arms were walled, and the other two arms were open. The maze was elevated 120 centimeters above the ground, and each arm measured 50 x 10 centimeters. The enclosed arms walls measured 30 centimeters high. The mice were placed in the maze’s center and allowed to move freely for 5 minutes. The total number of arm entries, the ratio of entries in open versus closed arms, distance traveled, and velocity were recorded and analyzed with Noldus EthoVision software.

### In vivo glutamate measurement (enzyme-coated microelectrode array)

Following behavioral testing, enzyme-immobilized amperometric microelectrode arrays (MEAs) were used to quantify tonic subregional glutamate levels and KCl-evoked glutamate release in hippocampal subregions, *dentate gyrus* (DG), *cornu ammonis 3* (CA3), and *cornu ammonis 1* (CA1), of anesthetized mice (n=6-12 mice per group), as we have previously described (Hunsberger *et al*. 2015c; Hunsberger *et al*. 2015a; Hunsberger *et al*. 2016b; Hunsberger *et al*. 2017).

#### Electrode preparation

Ceramic multisite microelectrodes arrays (MEAs) with eight paired platinum recording sites were purchased to measure glutamate and remove interferents (Quanteon, L.L.C. – Nicholasville, KY). The electrodes were prepared as previously described (Hunsberger *et al*. 2017; Burmeister & Gerhardt 2001). Briefly, sentinel sites were modified with an inactive protein matrix consisting of bovine serum albumin and glutaraldehyde and coated on the first pair of recording sites to block large molecule interferents. Then, a glutamate oxidase (GluOx) matrix was drop-casted on the first pair of recording sites to oxidize glutamate into alpha-ketoglutarate and hydrogen peroxide, the reporter molecule. A size exclusion layer was then electropolymerized onto the electrode to block large molecule interferents. Before implantation, the electrodes were calibrated, and sentinel sites were subtracted from the current obtained on the GluOx recording sites, producing a selective measure of extracellular glutamate. A silver/silver chloride (Ag/AgCl) reference electrode was used in both *in vitro* calibration and *in vivo* studies to provide a remote electrical reference point with electron flow to and from the working electrode.

#### Calibration

MEAs were calibrated before their use *in vivo* to create a standard curve for converting current to glutamate concentration and ensure glutamate sensitivity and selectivity. Using the Fast 16-MkII system (Quanteon, LLC), a constant potential of +0.7 V versus an Ag Ag/Cl reference (BASi, inc.) was applied to oxidize the reporter molecule, and the resultant current was amplified, digitized, and filtered. Calibration consisted of submerging the MEA tip into a well-stirred beaker of 40 mL 0.05 M phosphate-buffered saline and baselined for at least 15 minutes before adding three additions of 20 μL glutamate to achieve concentrations of 20, 40, and 60 μM. A slope was calculated to convert the increase in current (nA) produced by oxidation to the known glutamate concentration. Ascorbic acid (250 μM) and dopamine (2 μM) were added as interferents to ensure that the MEA maintained selectivity for glutamate. The limit of detection (LoD) was calculated and defined as the smallest concentration of glutamate that the electrode can measure, determined by a three-fold change over baseline noise or standard deviation.

#### Micropipette assembly

A glass micropipette (A-M systems) was pulled using a Narshige PC-10 and bumped to have an internal diameter of 10-15 μm. Successfully bumped pipettes were fixed to the microelectrode with the tip 75-100 μm from the reporter and sentinel sites with stick wax. The micropipette was then attached to a Picospritzer III (Parker-Hannifin, Cleveland, OH) and set to deliver consistent volumes of 100 nL at 0.138-1.38 bar pressure for 0.1-3.0 seconds. The spacing of the micropipette from the electrode was measured post-surgery to ensure that it did not shift during the procedure. Injection volume was quantified via displacement and was measured using a reticle-affixed stereoscope. The Picospritzer and micropipette were used to inject sterile-filtered isotonic 70 mM KCl. KCl solution was injected to measure evoked presynaptic release quanta at 100 nL/injection. Injections were performed with a Picospritzer III, and a stereomicroscope was used to measure volume displacement.

#### In vivo anesthetized glutamate recording

Surgical procedures were performed as described previously (Hunsberger *et al*. 2015c; Hunsberger *et al*. 2015a; Hunsberger *et al*. 2016b; Hunsberger *et al*. 2017). Briefly, mice were anesthetized with 4% isoflurane in an induction chamber and immediately placed in ear bars on a stereotaxic device (Kopf; Tujunga, CA). Mice remained on isoflurane ventilation (1-2.5%), and a self-regulating infrared heating pad and rectal probe were used to maintain the body temperature at ∼37°C to prevent anesthesia-induced hypothermia. When administered under controlled body temperature, isoflurane anesthesia does not alter resting glutamate concentrations or tau hyperphosphorylation (Tan *et al*. 2010). MEA recordings were taken in the CA1, CA3, and DG of the hippocampus, and stereotaxic coordinates were determined from bregma (Paxinos & Franklin 2012) [DG (AP: -2.3mm, ML: +/-1.5mm, DV: 2.1mm), CA1 (AP: -2.3mm, ML: +/-1.7mm, DV: 1.4mm), CA3 (AP: -2.3mm, ML: +/-2.7mm, DV: 2.25mm)]. MEA recordings were performed at a sampling rate of 10 Hz using constant potential amperometry (FAST-16 MkII). A miniature reference electrode was placed under the skin. Recordings were performed after reaching a stable baseline (20-45 minutes). Drug injections were performed in both hemispheres. Injection order was counterbalanced in hippocampal subregions and hemispheres. Data from some hippocampal subregions were excluded for reasons including clogging or failure of the MEA and death during surgery. Tonic glutamate, a measure of resting extrasynaptic glutamate, was calculated over 10 seconds in each subregion before injections. For KCl-evoked glutamate release, four to six injections of 100 nL of 70 mM KCl spaced 2-3 minutes apart were compared per group. Amperometric data were analyzed using a custom Matlab program.

#### Extracellular field recordings

Local field potential electrophysiology recordings were performed on the Schaeffer’s collateral pathway of the hippocampus, one week post behavioral testing as described previously (Pinky *et al*. 2023a). Animals assigned to electrophysiology (n=5-7 mice per group) were euthanized via CO_2_, and the brain was immediately removed, dissected, and placed into the continuously oxygenated artificial cerebrospinal fluid (aCSF, in mM; 124 NaCl, 2.5 KCl, 1.5 MgCl_2_, 2 CaCl_2_, 1.25 NaH_2_PO_4_, 25 NaHCO_3_, 25 dextrose, pH 7.4). Coronal brain slices were taken at 350 μm on a vibratome immersed in an ice-cold cutting solution and incubated in aCSF for a minimum of one hour at room temperature.

The slices were then transferred to a recording chamber and maintained under aCSF perfusion at 34°C bubbled with carbogen 95%CO_2_/5%O_2_ during recording. A bipolar stereotrode (Microprobes) was placed at the CA3 region, and a glass pipette filled with aCSF was placed at the stratum radiatum of the CA1 for recording field excitatory postsynaptic potentials (fEPSP) in the Schaeffer’s collateral pathway. To examine basal synaptic transmission, slopes of fEPSP were plotted in relation to stimulus intensities ranging from 0-200 μA at 20 μA intervals and represented as input-output response. Paired pulse facilitation (PPF) was measured by giving successive stimuli at varying time intervals, and the ratio of the second fEPSP (fEPSP_2_) to the first fEPSP (fEPSP_1_) was calculated and plotted. Long-term potentiation (LTP) was induced using a theta-burst stimulation (TBS) protocol after at least ten minutes of stable baseline. TBS consists of ten bursts of stimuli, each containing four pulses at 100 Hz, with an inter-burst interval of 200 milliseconds and 20 seconds between the five individual sweeps. Each slice was recorded for 60 min post-TBS, and LTP was calculated as the average of the fESPSs between minutes 50 and 60 post-TBS induction. Data was recorded and analyzed with WinLTP software (University of Bristol, UK).

#### Immunohistochemistry

Mice were transaortically perfused with ice-cold phosphate-buffered saline (PBS) following ketamine/xylazine anesthesia (n=3 mice per group). The brain was dissected and immersion fixed in 4% paraformaldehyde at 4°C for a minimum of 24 h. The samples were then sequentially transferred into 10%, 20%, and 30% sucrose solutions in PBS and stored at 4°C for a minimum of 24 h per solution. Excess moisture was removed from the tissue exterior, and tissue was flash-frozen in dry ice and stored at -80°C. Tissues were then mounted in a Leica CM3050S cryostat using Optimal Cutting Temperature compound (OCT, Sakura Tissue-Tek), and 40 μm coronal hippocampal slices were taken and placed in a 0.1 M phosphate-buffered saline (PBS). Sample sections were then permeabilized with 0.5% Triton-X 100 in PBS (Fisher Scientific, Waltham, MA) for ten minutes and then blocked in 5% goat serum for 20 minutes. Tissue was incubated overnight in primary antibody vGLUT1 (cat. no. 135-303, Synaptic Systems) or 6E10 (cat. no. 9347-02, Biolegend). Sections were washed in PBS and incubated in fluorophore-conjugated secondary antibodies for two hours in the dark. Sections were again washed and mounted with Permount (Fisher Scientific) between a gelatin-coated microscope slide and coverslip. Unless otherwise noted, all incubation and wash steps were performed on a shaker at 40 rpm at room temperature.

Fluorescence imaging was performed with a Nikon Eclipse Ti2 and a Nikon A1 confocal microscope. Images were analyzed using ImageJ by delineating appropriate subregions and splitting the channels. All images underwent postprocessing consisting of a 50-pixel rolling ball background subtraction. Masks were created to identify individual pixels with intensity of interest (VGlut1, Synaptophysin, 6e10) by thresholding images to select the top 25 ± 1% counts per pixel. Further, different regions of interest were identified in each IHC image that represented different regions within the hippocampus. Quantified intensities were then separated by region and compared. Individual pixel intensities were then integrated using custom-written Matlab scripts (R2022b). The ratio or absolute value of intensities was quantified on a pixel-by-pixel basis within each region of interest. Finally, mean values for each region were quantified by cumulative distribution fitting curve and aggregated as final results. Statistical comparisons were performed on the cumulative distributions.

#### ELISA

The Aβ_1-42_ concentrations were quantified using values obtained from a standard curve associated with the ELISA kit following the manufacturer’s protocol (cat. no. DAB142; R&D Systems, Minneapolis, MN, USA). The optical densities of each well were measured at 450 nm using a microplate reader (Agilent, Santa Clara, CA, USA), and the sample Aβ_1–42_ concentrations were determined by comparison with the Aβ_1–42_ standard curves (n=4-6 mice per group). All readings were in the linear range of the assay. Values were standardized to total protein concentrations.

#### Western Blot

Mouse hippocampal tissue was homogenized in radioimmunoprecipitation assay buffer (50mM Tris-HCl pH 7.4, 150mM NaCl, 0.5% Triton X, 1mM EDTA, 3% SDS, 1% deoxycholate) and protease/phosphatase inhibitor cocktail (5872S, Cell Signaling, Danvers, Massachusetts, USA), then samples were centrifuged at 4°C and a speed of 13,000 rpm for 90 minutes. Hippocampal samples (n=3-4 mice per group) were stored at -80°C. A bicinchoninic acid (23227; Thermo Fisher Scientific, Waltham, MA, USA) protein assay using bovine serum albumin (23209; Thermo Fisher Scientific, Waltham, MA, USA) as a standard was used to determine protein concentrations. Samples were centrifuged a second time at 4°C at a speed of 13,000 rpm for 90 minutes prior to performing the protein quantification. 15 ug of sample protein was added 4X Laemmli Buffer (1610747; BioRad, Hercules, CA, USA), and samples were heated for 5 min to 95 °C then loaded on 10% hand-cast sodium dodecyl sulfate page gels. Gels were then transferred overnight onto LF PVDF 0.45um membrane (1620264; BioRad, Hercules, CA, USA). After transfer, membranes were blocked under gentle agitation for 1 h at room temperature (∼23°C) with equal parts Intercept PBS blocking buffer (927-70001; Li-Cor, Lincoln, NE, USA) and 1M PBS; this blocking buffer mixture was also used as primary antibody diluent for incubation. Membranes were then incubated with primary antibodies directed against the protein of interest overnight (16 hours) at 4°C. Primary antibodies include conjugated Tau13 Alexa Flour 680 (1:5000) (sc-21796 AF680, Santa Cruz Biotechnology, Inc. Dallas, TX, USA) and phospho-Tau Ser396/Ser404 (PHF-1) (1:5000) (sourced from Dr. Peter Davies). Following five 5-minute washes of the membrane with TBST (20 mM Tris, 150 mM NaCl, 0.1% Tween 20, pH 7.4), an IgG secondary antibody IRDye 800CW Goat anti-mouse (1:3000) (926-32210; Li-Cor, Lincoln, NE, USA) diluted in wash buffer was added for a 1-hour room temperature incubation with gentle agitation. Immediately following the incubation, the five washes were repeated with TBST and then stored temporarily in TBS until NIR imaging.

#### Statistics

Statistical analysis was performed using JMP 16.0 (Sas, Cary, NC), FIJI, and Matlab R2022b), and all graphs were prepared using GraphPad or Biorender.com. Statistical procedures include two-sided t-tests, ANOVAs, or repeated measures ANOVA (RMANOVA), two-sided Kolmogorov-Smirnov (KS) tests, Mann-Whitney U tests, and Tukey *post hoc* comparisons were performed if a significant omnibus test was observed. The critical alpha level was set to 0.05, and numerical results are presented as means ± SEM.

## Results

### 3xTg neurons express increased VGlut1 and number of presynaptic SVs compared to TgNeg

To determine if presynaptic VGLUT1 and vesicle release are increased in 3xTg, we used the pH-sensitive VGlut1-pHluorin approach in hippocampal cell cultures (Maschi *et al*. 2021). Neurons were transfected with VGlut1-pHluorin which is dark when SVs are in the recycling pool and becomes bright when SVs are exposed to the pH-neutral extracellular environment during exocytosis (**Fig. 1A**). We determined presynaptic SV release mechanics by quantifying VGlut1- pHluorin intensity at the single presynapse level using the following approach: we first imaged presynapses in the absence of stimulation to measure spontaneous release frequency (Leitz & Kavalali 2014); (ii) we then stimulated neurons at 1 Hz to measure release probability (Maschi *et al*. 2021); (iii) we then stimulated neurons at 40 Hz to measure the size of the recycling pool (Taipala *et al*. 2022); (iv) and finally we exposed neurons to NH_4_Cl to measure the total SV pool size (Taipala *et al*. 2022). These measurements were performed sequentially to determine changes at the single presynaptic level (**Fig. 1B**).

**Figure 1.**
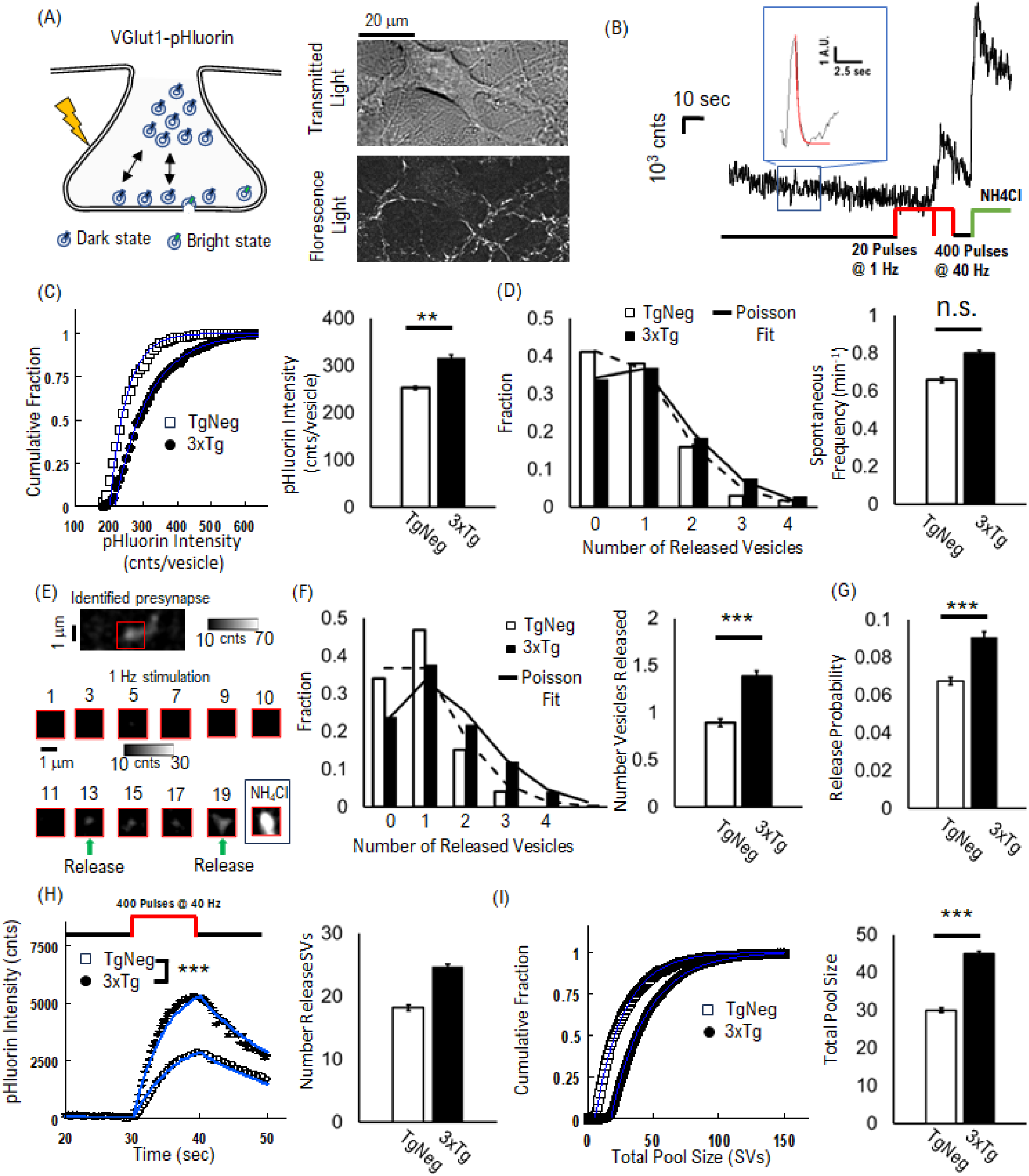
3xTg cultured hippocampal neurons express increased VGlut1 and vesicle release mechanics. (A) Representation of VGlut1-pHluorin fluorescent protein when dark within the acidic environment of the SV, and bright during exocytosis. Example of raw VGlut1-pHluorin intensity during presynaptic activity. (B) Representative trace of VGlut1-pHluorin intensity integrated for a single presynapse. A single spontaneous SV release trace is shown (inset). Presynapses are imaged in the absence of stimulation for 80 sec, followed by 1 Hz electrical stimulation for 20 pulses, then 40 Hz electrical stimulation for 400 pulses, and finally, exposure to NH_4_Cl. (C) Cumulative distribution of single SV VGlut1-pHluorin intensities. Peak intensity from single SV release events (inset, (B)). Mean VGlut1-pHluorin intensity per SV bar plot (right panel) from fits to cumulative data (Blue lines, left panel). (D) Spontaneous release distributions (bars) from identified single SV release events fit to Poisson distribution (solid lines). Mean spontaneous release frequency per min (Right panel) obtained from Poisson distribution fits. (E) Example of raw single SV release events during 1Hz stimulation. (F) Distribution of release events during 1Hz stimulation (Left panel). Data fit to Poisson distribution (solid lines). Mean number of released SVs (Right panel) from Poisson distribution fits (Solid lines, left panel). (G) Calculated release probability calculated as number of released SVs divided by total number of pulses. (H) Integrated VGlut1-pHluorin intensity during 40Hz stimulation (left panel) and computational model fits (solid lines). Total number of released SVs during 40Hz stimulation (Right panel) calculated from computational model fits (solid lines, left panel). (I) Cumulative distribution of maximum VGlut1-pHluorin intensity during NH4Cl exposure. Intensity normalized to mean single SV intensity (bar plot in (C)) resulting in expected number of SVs per presynapse. Mean total pool size (Right panel) obtained from cumulative fits (solid lines, Left panel). Error bars from cumulative fits are SEM from normal distributions. N_SV_ CT = 237; 3xTg = 270 N_presynapses_ CT = 400; 3xTg = 401 N_samples_ CT = 2; 3xTg = 2 N_cultures_ CT = 1; 3xTg = 1 Two-tailed t-Test comparisons were performed for (D),(F),(G). Two-tailed KS-Test comparisons were performed for (C),(I). Mann-Whitney U test performed for (H). *** = p<.001; ** = p<.01

We first determined the change in VGlut1 per SV by comparing intensities from single SV spontaneous release events in the absence of stimulation (**Fig. 1C**). We observed that 3xTg VGlut1-pHluorin intensity distributions exhibited the same variance but with a statistically significant larger average than TgNeg (**Fig. 1C**). This increase in 3xTg is equivalent to a factor of ∼1.2 compared to TgNegs, which is lower than the ∼1.4 increase in VGlut1 we have previously measured in the rTg(P301L)4510 mouse model (Taipala *et al*. 2022). However, a 1.2 VGlut1- pHluorin intensity increase supports an integer increase in the number of VGlut1 transporters per SV we previously proposed (Taipala *et al*. 2022), with a baseline of 5 +/- 1 VGlut1 transporters per SV for TgNeg and an increase to 6 +/- 1 VGlut1 transporters per SV for 3xTg.

We then quantified spontaneous release frequency to determine if baseline presynapse release is increased along with the number of VGlut1 transporters (**Fig. 1D**). Here, we counted the number of single SV release events in the absence of stimulation using a previously established approach (Leitz & Kavalali 2014), and then fit the distribution of SV release events to a Poisson distribution (see methods). We found that both 3xTg and TgNeg presynapses exhibit the same spontaneous release frequency consistent with previously measured frequency in hippocampal neurons (Leitz & Kavalali 2014); however, spontaneous release was elevated by a factor of ∼1.3 for 3xTg presynapses compared to TgNeg (**Fig. 1D**). These results support the hypothesis that both glutamate release amount and frequency is increased in 3xTg mice compared to TgNeg littermates.

Along with increased spontaneous release, we sought to establish if baseline release probability is increased in 3xTg neurons. We quantified release probability as the number of SV release events measured during 1 Hz stimulation (**Fig. 1B**). Hippocampal presynapses have a low release probability (<= 0.1) (Gramlich & Klyachko 2019), resulting in the majority of presynapses releasing one or no SVs during stimulation (**Fig. 1E**). We then fit the distribution of release events per synapse and observed an ∼1.4x increase in the number of released vesicles for 3xTg neurons compared to TgNeg (**Left Panel, Fig. 1F**). We then divided the average number of release events by the number of stimulus pulses (20 pulses, **Fig. 1B**) resulting in a ∼1.3x increase for 3xTg presynapses compared to TgNeg (3xTg: 0.083; TgNeg: 0.064; **Fig. 1F**).

To determine if increased release probability supports an increase in sustained release, we quantified VGlut1-pHluorin intensity during a bout of extended stimulation at the individual presynapse level (40 Hz., **Fig. 1B**). We observed a significant increase in average VGlut1- pHluorin intensity for 3xTg neurons compared to TgNeg neurons (**Fig. 1G**), consistent with our previous observations in the rTg(P301L)4510 mouse model (Taipala *et al*. 2022). We then fit the VGlut1-pHluorin intensity (**Blue Lines, Fig. 1G**) using our previously established computational model (Taipala *et al*. 2022), using the experimentally measured VGlut1-pHluorin intensity per vesicle (**Fig. 1C**) and release probability (**Fig. 1F**). The model fits showed a ∼1.4x increase in the number of released vesicles for 3xTg neurons compared to TgNeg (3xTg: 25; TgNeg: 18; **Fig. 1G**).

Finally, we sought to establish if the total SV pool size was altered to support increased spontaneous and stimulated SV release. To determine total pool size, we quantified the maximum VGlut1-pHluorin intensity during exposure to NH_4_Cl (NH_4_Cl **Fig. 1B**), divided by the average intensity per single SV (**Fig. 1C**), which is equivalent to the number of SVs within a presynapse. We observed both 3xTg and TgNeg neurons exhibit pool sizes similar to previous studies of hippocampal neurons (Balaji & Ryan 2007). However, we also observed a significant increase in the distribution of SV pool sizes for 3xTg neurons compared to TgNeg neurons (**Fig. 1H**). We then fit the distribution (Blue Lines, **Fig. 1H**) to determine the average pool size. We observed a ∼1.5x increase in the number of vesicles for 3xTg neurons compared to TgNeg (3xTg: 45; TgNeg: 30; **Fig. 1H**). This is consistent with previous studies showing an increase in SV pool size for Aβ mutation (Hark *et al*. 2021), suggesting differential roles for tau and Aβ that cumulatively increase glutamate release: where tau increases VGlut1 levels and Aβ increases both VGlut levels and SV pool size.

These combined VGlut1-pHluorin results show that 3xTg neurons exhibit increased VGlut1 expression per vesicle, an increase in presynaptic release mechanics (spontaneous and stimulated release probability), as well as an increase in total pool size. These combined changes support the hypothesis that the combined 3xTg mutations mediate hyperexcitability.

#### VGlut1 levels are increased in 3xTg mice and decreased following treatment with troriluzole

Increased *in vitro* VGlut1 expression in cell cultures does not guarantee that VGlut1 expression is elevated in the hippocampus *in vivo* or that it can be rescued. To support that VGlut1 levels remain elevated during disease progression and can be rescued *in vivo* with troriluzole, we measured relative VGlut1 expression in hippocampal slices (**Fig. 2**). We measured VGlut1 expression relative to the synaptic vesicle protein, synaptophysin (Syn), using immunohistochemistry in 8-month-old 3xTg and negative control littermates (TgNeg), as well as 3xTg mice treated with troriluzole for 3 months.

**Figure 2.**
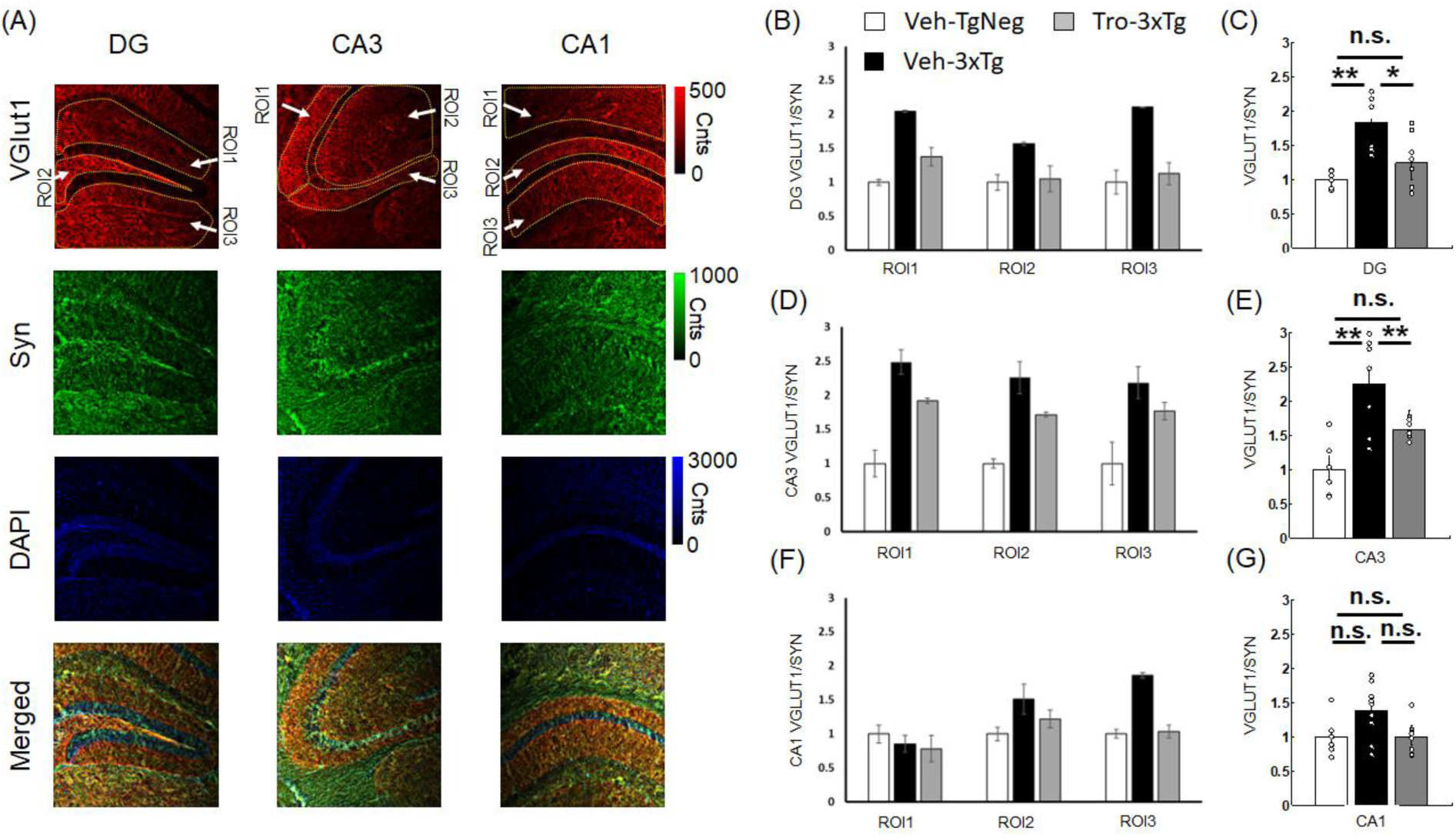
Troriluzole decreases the elevated VGlut1 levels observed in 3xTg mice. (A) Example images of slices stained for VGlut1 (Red), Syn (Green), DAPI (Blue), and merged (Bottom). Within each area, three distinct regions of interest (ROI1, ROI2, ROI3) are identified (Dotted outlines) and the VGlut1/Syn intensity ratio quantified separately for each ROI or aggregated across the regions for the DG (B & C), CA3 (D & E), and CA1 (F & G). All means normalized to TgNeg. Error bars are SEM across all regions of the same group (DG, CA3, CA1). * = p<.05, ** = p<.01, *** =p<.001. All statistics from two-tailed KS-test.

We first separated IHC hippocampal images across three different areas and three regions of interest within each area (**Fig. 2A**). We imaged the dentate gyrus (DG, **Fig. 2A**) in regions representing overlap with the Schaffer collateral (ROI1), inputs from granular cells (ROI2), and mossy fiber outputs to the CA3 cells (ROI3). We imaged the Cornu Ammonis 3 (CA3, **Fig. 2A**) in regions representing inputs from the medial septum (ROI1) (Salib *et al*. 2019), the Schaffer Collateral (ROI2) (Cherubini & Miles 2015; Basu & Siegelbaum 2015), and mossy fiber inputs from the DG cells (ROI3) (Basu & Siegelbaum 2015). Finally, the Cornu Ammonis 1 (CA1, **Fig. 2A**) was separated into regions representing the cortex outside the hippocampus (ROI1) (Swanson & Hahn 2020), the Schaffer Collateral (ROI2) (Basu & Siegelbaum 2015), and a mixture of temporoammonic path and perforant path inputs from the entorhinal cortex (ROI3) (Basu & Siegelbaum 2015).

The quantified distribution of relative VGlut1/Syn expression in each region showed an increase in VGlut1 expression for all hippocampal regions and areas for 3xTg mice (Black, **Fig. 2 B, D, F**) relative to TgNeg mice (White, **Fig. 2 B, D, F**). Overall, 3xTg VGlut1 expression was between 1.5x (CA1, Fig. 2 G) and 2.2x (CA3, **Fig. 2 E**) higher than TgNeg mice. The only region where VGlut1 expression was unchanged in 3xTg mice was outside the hippocampus (ROI1 in CA1, **Fig. 2F**). These results show that increased VGlut1 expression per synaptic vesicle observed in cell cultures (**Fig. 1**) occur *in vivo* in 8-month-old mice.

We then quantified VGlut1 expression for 3xTg mice exposed to troriluzole (Grey, **Fig. 2 B, D, F**). We observed a significant reduction in VGlut1 expression in all regions across all hippocampal areas. In particular, VGlut1 expression decreased to the same amount as TgNeg mice in both the DG (**Fig. 2C**) and CA1 areas (**Fig. 2G**). In the CA3 area, VGlut1 expression in troriluzole-treated 3xTg was lower than that of vehicle-treated 3xTg mice but remained higher than TgNeg mice (**Fig. 2E**). These results show that troriluzole reduces VGlut1 expression in 3xTg mice.

#### Troriluzole rescues the elevated hippocampal tonic glutamate and KCl-evoked glutamate release observed in 3xTg mice

We used microelectrode arrays to determine whether 3xTg mice exhibit increased tonic and KCl-evoked release of glutamate similar to what we previously observed in the rTg(P301L)4510 mouse model (Hunsberger *et al*. 2015b; Hunsberger *et al*. 2015c; Hunsberger *et al*. 2021) and others have seen in APP/PS1 mice (Hascup *et al*. 2021; Hascup & Hascup 2015; Hascup *et al*. 2019). We also sought to determine whether troriluzole could restore observed alterations to that of controls. Veh-3xTg mice exhibited increased tonic glutamate levels in all three hippocampal subregions, an effect attenuated by troriluzole treatment (DG: *F*_(2,21)_ = 11.6, *p* = .0004; CA3: *F*_(2,23)_ = 6.7, *p* = .0055; CA1: *F*_(2,26)_ = 16.7, *p* <.0001; **Fig. 3A**). The capacity for glutamate release was examined by local application of 100 nL of 70 mM KCl via a micropipette, which produced reproducible glutamate release (**Fig. 3B**). The amplitudes of KCL-evoked glutamate release differed significantly amongst the groups in all three subregions (DG: *F*_(2,172)_ = 9.2, *p* = .0002; CA1: *F*_(2,175)_ = 49.8, *p* < .0001; and CA3: *F*_(2,213)_ *=* 36.8*, p* < .0001; **Fig. 3C**); 3xTg mice exhibited significant increases in evoked release that were rescued by treatment with troriluzole treatment (**Fig. 3B,C**).

**Figure 3.**
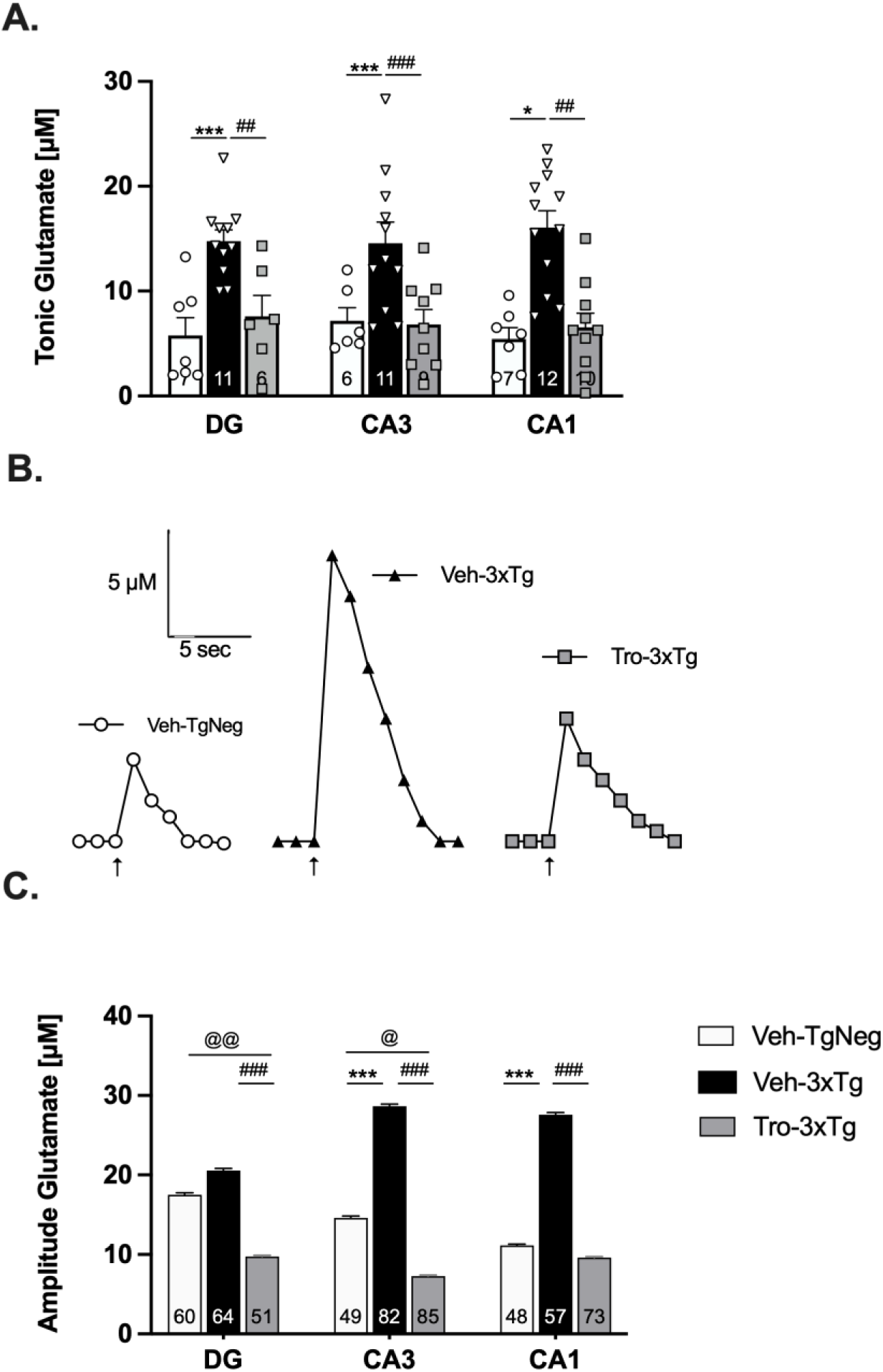
Extracellular tonic and potassium chloride (KCl)-evoked release of glutamate in the DG, CA3, and CA1 regions of the hippocampus. (A) Tonic glutamate levels were significantly increased in Veh-3xTg mice in all hippocampal subregions, an effect attenuated by troriluzole treatment. n = 6-12 mice per group. (B) Representative recordings of baseline-matched KCl-evoked glutamate release in the CA1 indicate that troriluzole treatment reduces the significant increase in the amplitude of glutamate release in Veh-3xTg mice. Local application of KCl (↑) produced a robust increase in extracellular glutamate that rapidly returned to tonic levels. (C) KCl-evoked glutamate release in Veh-3xTg mice was significantly increased after local application of KCl and was attenuated with troriluzole treatment. n = 48 – 82 KCl injections per group from 6-12 mice per group. * represents significant differences between Veh-TgNeg and Veh-3xTg; # represents significant difference between Veh-3xTg and Tro-3xTg; @ represents significant differences between Veh-TgNeg and Tro-3xTg, */#/@ *p* < .05, **/##/@@ *p* < .01, ***/###/@@@ *p* < .001

#### Troriluzole restores the basal synaptic transmission and synaptic plasticity deficits of 3xTg mice

To determine whether troriluzole rescues deficits in glutamatergic basal synaptic transmission in the Schaffer collateral pathway of the hippocampus in 3xTg-AD mice, fEPSPs were measured at increasing stimulus intensities. fEPSP slope was reduced in Veh-3xTg compared to Veh-TgNeg mice, indicating deficits in baseline glutamatergic synaptic transmission. Treatment with troriluzole increased the fEPSP slope of 3xTg mice (*F*_(2,18)_ = 6.7, *p* = .008; **Fig. 4A**). To determine whether alterations in basal synaptic transmission could be related to alterations in presynaptic glutamate release, we evaluated paired-pulse facilitation (PPF), a type of short-term plasticity that depends on residual calcium build-up in the presynaptic terminal (Zucker 1989). We observed a decrease in PPF in Veh-3xTg mice that was attenuated by troriluzole treatment (*F*_(2,18)_ = 5.4, *p* = .01; **Fig. 4B**). Because there is an inverse relationship between PPF and presynaptic release probability, the decreased PPF value of Veh-3xTg mice indicates that 3xTg mice have increased presynaptic release probability, similar to what was observed in cultures (**Fig. 1**), and this increase was reduced by treatment with troriluzole.

**Figure 4.**
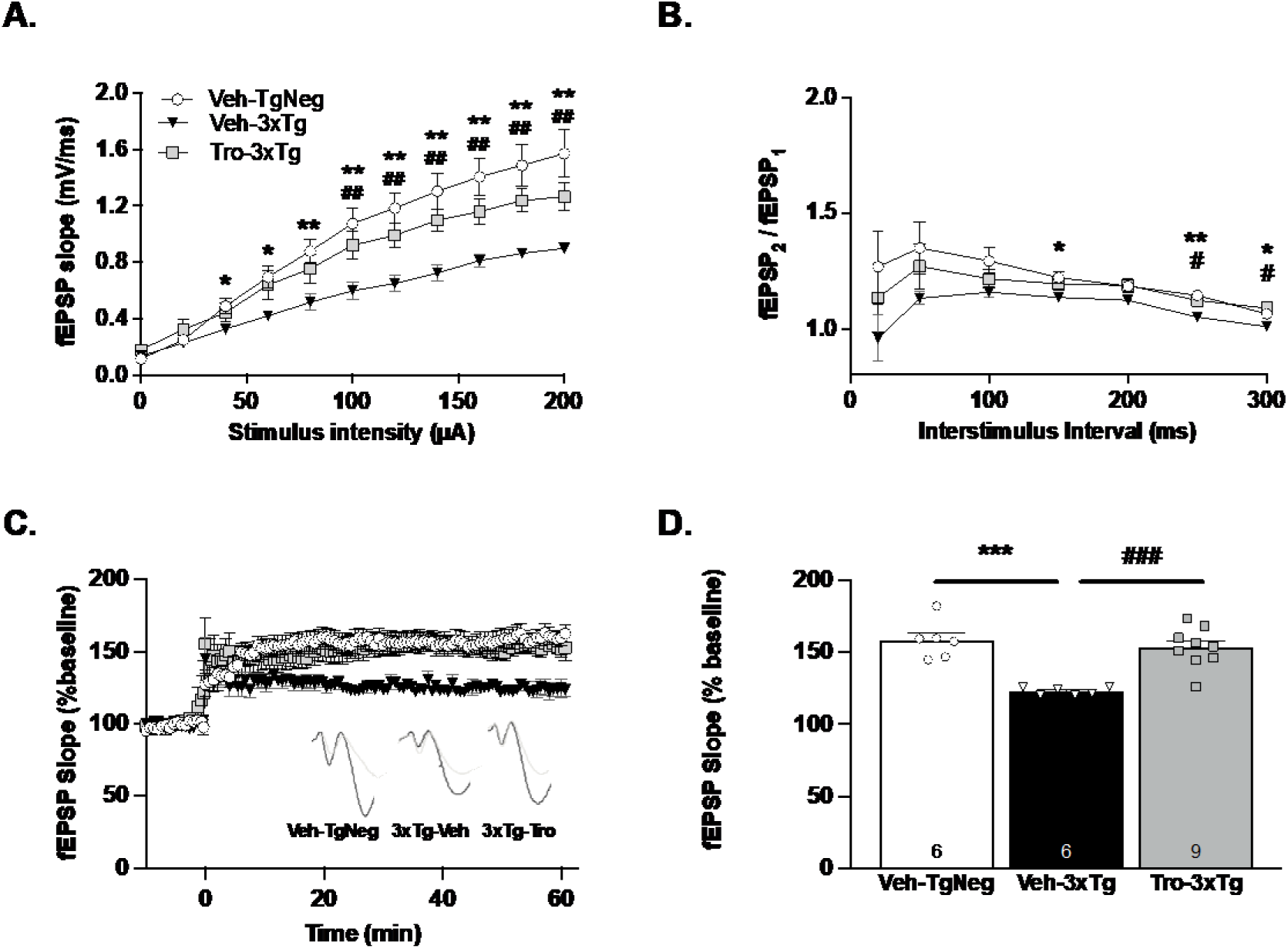
Troriluzole treatment rescues hippocampal field electrophysiology alterations in 3xTg mice. (**A**) Input-output curve of fEPSP slope measured at increasing stimulus intensities. Troriluzole rescued deficits in basal synaptic transmission in 3xTg mice. (**B**) Paired-pulse facilitation (PPF) expressed as the ratio of the second stimulus fEPSP slope to the first stimulus fEPSP slope plotted as a function of interstimulus interval. Troriluzole rescued alterations in PPF in 3xTg mice. (**C)** Long-term potentiation (LTP) graph represents fEPSP slope before and after induction by theta-burst stimulation (TBS). (**D**) LTP bar graph shows the average of fEPSPs recorded during the time period 50-60 min following TBS induction normalized to baseline levels. Troriluzole rescues deficits observed in LTP in the 3xTg mice. * represents significant differences between Veh-TgNeg and Veh-3xTg; # represents significant difference between Veh-3xTg and Tro-3xTg; */# *p* < .05, **/## *p* < .01, ***/### *p* < .001; n = 6-9 slices from 5-7 mice per group.

We next measured LTP, the cellular correlate of learning and memory (Luscher & Malenka 2012), to determine whether basal synaptic transmission alterations affected synaptic plasticity and whether troriluzole could rescue these alterations. 3xTg-Veh mice exhibited deficits in LTP that were rescued by troriluzole administration (*F*_(2,18)_ = 16.5, *p* < .0001; **Fig. 4C,D**).

#### Troriluzole decreases hippocampal Aβ load and tau levels

We next sought to examine Aβ load detected by 6E10 antibody in 3xTg mouse brains as determined by immunofluorescence analysis (**Fig. 5A**). While no differences were observed between 3xTg mice treated with vehicle versus troriluzole for the DG (p = 1, two-tailed KS-test; **Fig. 5B)** or CA3 (p = 0.76, two-tailed KS-test; **Fig. 5C**), troriluzole treatment significantly reduced Aβ load for the CA1 region (p = 0.0028, two-tailed KS-test; **Fig. 5D**). Troriluzole treatment also significantly reduced hippocampal Aβ_42_ levels in 3xTg mice as assessed using ELISA (p=.0036; two-tailed t-test; **Fig. 5E**). Troriluzole reduced tau levels detected by the Tau-13 antibody specific for human tau (p=.013; two-tailed t-test) and marginally decreased tau phosphorylation at Ser396/404, as detected with the PHF1 antibody (p=.1; two-tailed t-test; **Fig. 5F,G**).

**Figure 5.**
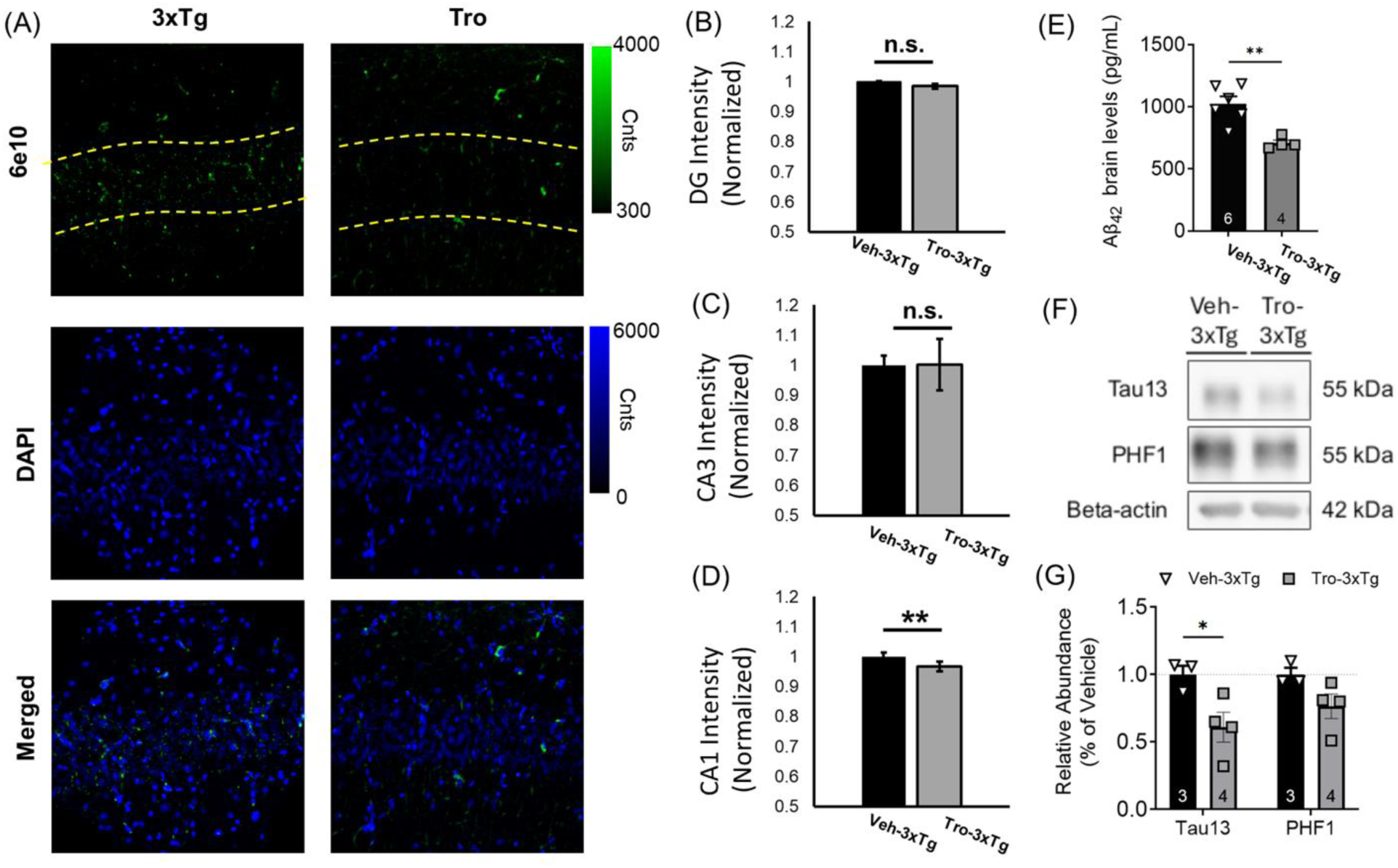
Troriluzole treatment reduces Aβ and tau. Example images of slices stained for 6e10 (Green), DAPI (Blue), and merged (Bottom). An example cornu ammonis 1 region is shown for TgNeg, 3xTg, and Troriluzole. The hippocampus was separately imaged in the dentate gyrus (DG), cornu ammonis 3 (CA3), and cornu ammonis 1 (CA1). The quantified mean 6e10 intensity ratio obtained by aggregating intensities per pixel for the same region in all samples of the same condition is shown for the DG (B), CA3 (C), and CA1 (D). Error bars are SEM of sample-to-sample variations, and a two-tailed KS-Test comparison was performed for all comparisons in B-D. n=3 mice per group. (E) Hippocampal Aβ_42_ levels in 3xTg mice as assessed using ELISA. n= 4-6 mice per group. (F) Representative images for Tau13, PHF1, and beta-actin. (G) Tau-13/beta-actin and PHF-1/beta-actin relative ratios from hippocampal lysate. n=3-4 mice per group. *** = P<0.001; ** = P<0.01

#### Troriluzole restores open/closed arm entry in EPM

3xTg mice have been shown to exhibit an anxiety-like phenotype in the elevated plus maze (Zhang *et al*. 2016). To assess the effect of troriluzole on anxiety-like behavior, the elevated plus maze (EPM) was used. No significant differences were observed in the distance traveled, average speed, or the average number of elevated plus-maze arm entries (*p*s > .05, **Fig. 6A-C**), indicating no difference in locomotor activity among groups. However, vehicle-treated 3xTg mice exhibited a reduction in the ratio of entries into open versus closed arms, suggesting an anxiety-like phenotype restored by treatment with troriluzole (*F*_(2,45)_ = 3.4, *p* = .04; **Fig. 6D**).

**Figure 6.**
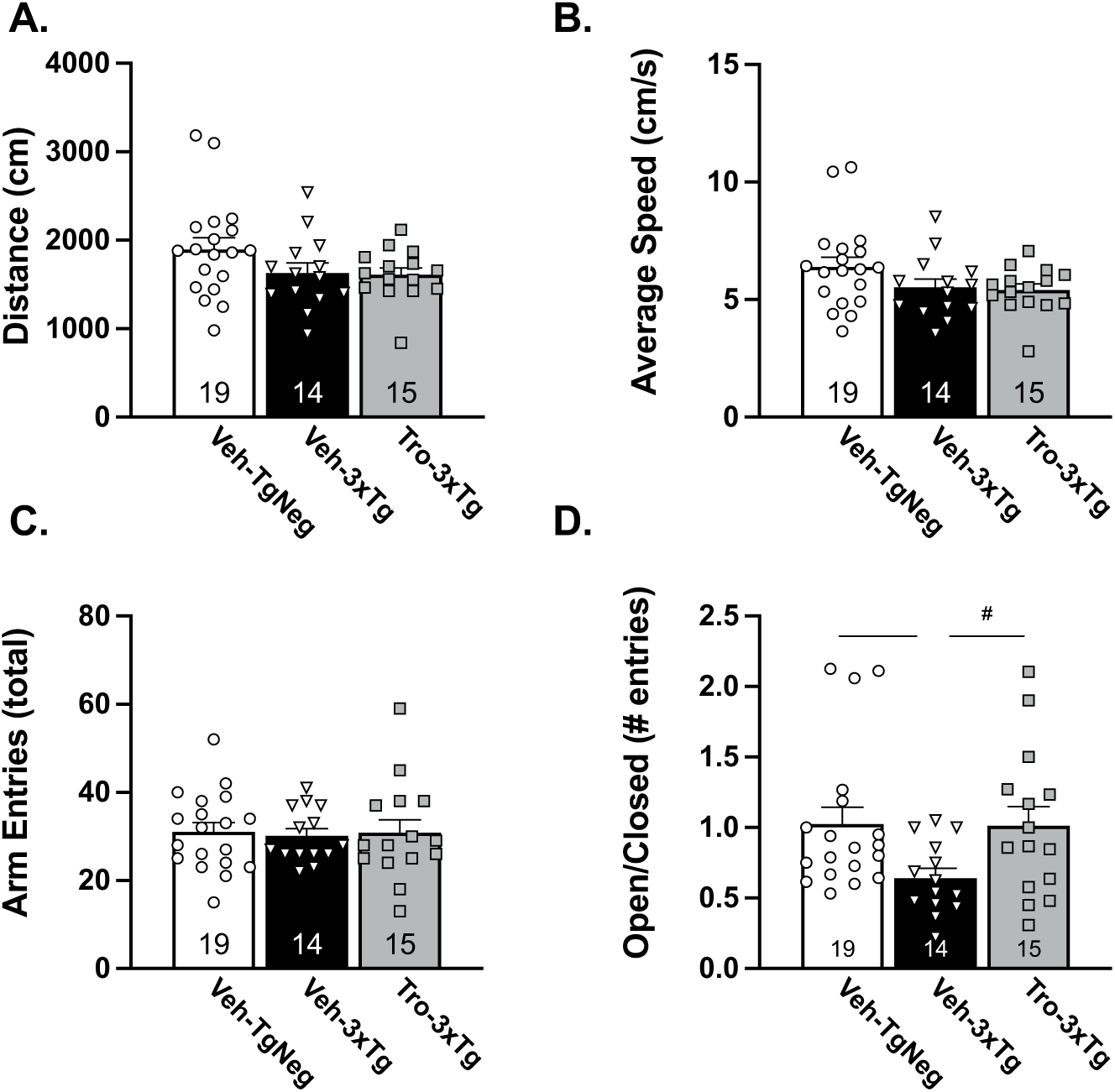
Troriluzole rescues the performance of 3xTg mice in an elevated plus maze. (**A-C**) Distance, average speed, and number of arm entries were similar among groups. (**D**) Troriluzole rescued the decrease in the open/closed arm entry ratio of 3xTg mice. (* represents significant differences between Veh-TgNeg and Veh-3xTg, # represents significant difference between Veh-3xTg and Tro-3xTg; */# *p* < 0.05, n = 14-19 mice per group).

#### Troriluzole rescues spatial reference memory deficits in the Morris water maze

We first compared the groups in the visible platform task to determine whether visual or motor functioning differed among the groups. As previously reported (Roda *et al*. 2020), swim speed was significantly increased in the vehicle-treated 3xTg group (*F*_(2,45)_ = 5.5, *p* = .008; **Fig. 7A**). However, pathlength in the visible platform task did not differ among the groups (*p* > .05; **Fig. 7B**), suggesting that neither transgene nor treatment impaired visual or motor functioning. We next compared spatial reference learning and memory using hidden platform testing with probe trials inserted on days 4-6 and 8. The groups did not differ in average path length during hidden platform trials (*p* > .05; **Fig. 7C**), though Veh-TgNeg (*F*_(5,14)_ = 6.5, *p*=.003) and Tro-3xTg (*F*_(5,9)_ = 5.6, *p*=.01) mice exhibited improved performance across trials that the Veh- 3xTg mice did not (*F*_(5,10)_ = 1.4, *p*=.31), indicating deficits in acquisition for the 3xTg mice that were rescued by troriluzole treatment. For the probe trials, vehicle-treated 3xTg mice, which exhibited chance performance (25% target quadrant occupancy), spent significantly less time in the target quadrant compared to both TgNegs and 3xTg mice treated with troriluzole (*F*_(2,45)_ = 37.5, *p* < .0001; **Fig. 7D-H**). Together, these data indicate that troriluzole treatment restores spatial reference memory in 3xTg mice.

**Figure 7.**
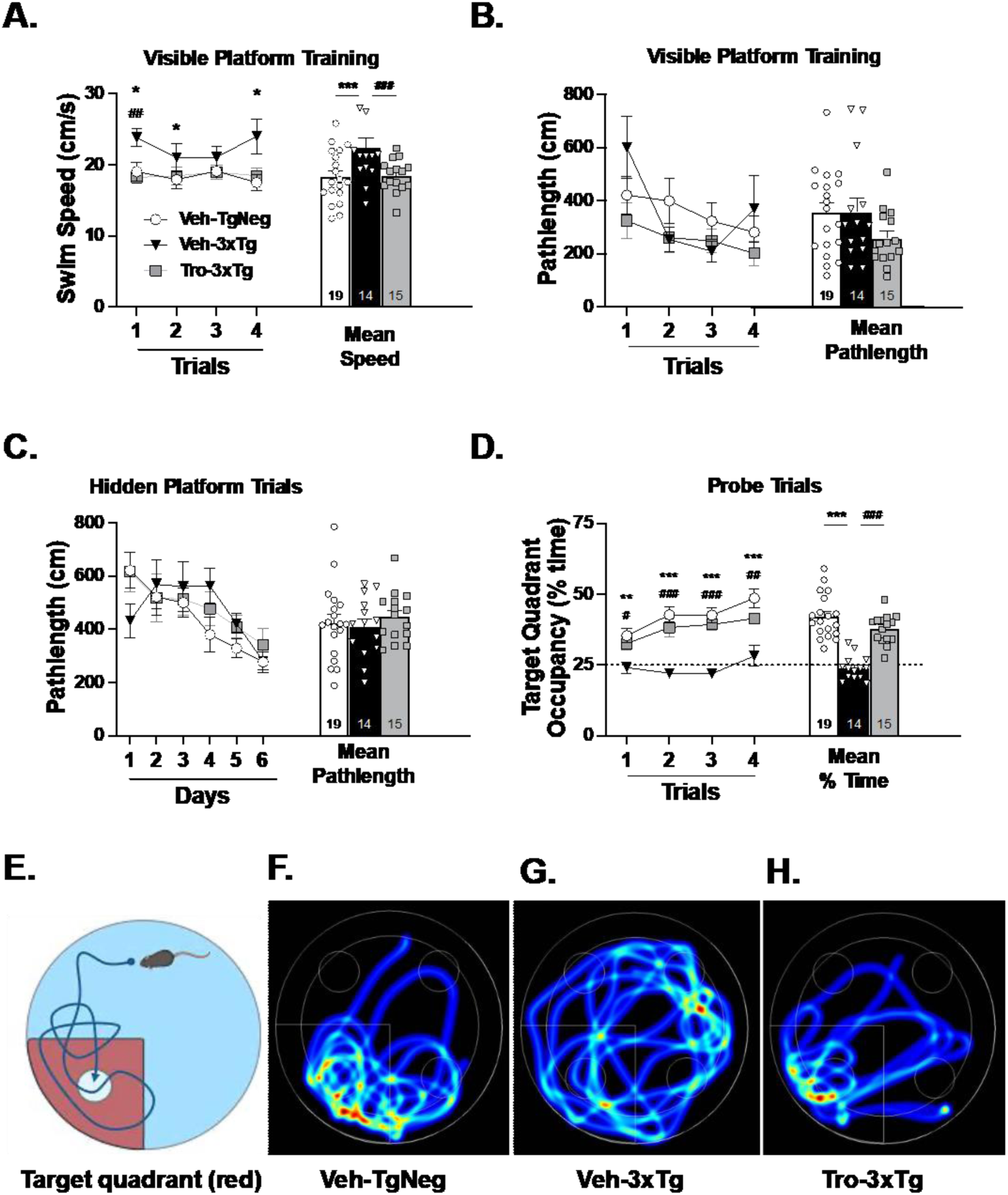
Troriluzole rescues deficits in reference memory observed in the Morris water maze. (**A**) Veh-3xTg exhibited an increased swim speed in the visual platform test. (**B**) No differences were observed in the pathlength to find the platform during the visual platform training trials. (**C**) Similar average pathlengths were observed amongst the groups. (**D**) Veh-3xTg mice exhibited a decreased time spent in the target quadrant during probe trials that were restored to TgNeg levels with troriluzole treatment. (**E**) Animation represents the target quadrant, or the quadrant in which the rescue platform was located, in hidden platform trials. The smaller circle represents the platform location. (**F-H**) Representative heatmap visualization of spatial occupancy during probe trials. The warmer the color correlates with an increased time spent in that location. * represents significant differences between Veh-TgNeg and Veh-3xTg; # represents significant difference between Veh-3xTg and Tro-3xTg; */# *p* < .05, **/## *p* < .01, ***/### *p* < .001; n = 14-19 mice per group.

## Discussion

The cell culture pHluorin approach used in the present study provides new details about molecular-level presynaptic function. Previous studies have shown that P301L tau induces increased VGlut1 expression (Hunsberger *et al*. 2021; Hunsberger *et al*. 2015c; Hunsberger *et al*. 2015a; Taipala *et al*. 2022), while other studies have shown that APP mutations lead to increased vesicle pool size in the presynapse (Hark *et al*. 2021). Here, we show that both occur simultaneously in the 3xTg mouse model (**Fig. 1**). Further, the present study shows that increased VGlut1 and increased synaptic vesicle number combine and lead to greater glutamate release (**Fig. 1**). The increase in VGlut1 expression and synaptic vesicle number also represents a significant increase in basal energy demand. Up to 45 percent of all presynaptic level energy demands go to maintaining glutamate concentration per vesicle under resting conditions (Pulido & Ryan 2021). Thus, increasing the number of VGlut1 transporters (which are ATP-dependent) and the number of vesicles will significantly increase the basal energy demands for glutamatergic synapses. Based on the results in the present study, hippocampal excitatory neuronal energy demand likely doubles at baseline from the combination of a 1.3 increase from VGlut1 (**Fig.1 C**) and a 1.5 increase from total pool size (**Fig.1 I**). This increase in demand would be further exacerbated during hyperexcitability when vesicles refill with glutamate after exocytosis. Thus, the molecular level results have significant implications on mitochondrial ATP production, which warrants further study. Overall, these presynaptic changes represent a significant molecular-level factor contributing to hyperexcitability. To further support this molecular-to-circuit level hypothesis, future studies should explore how preventing the VGlut1-mediated increased glutamate release, in turn, rescues network-level hyperexcitability.

The VGlut1 IHC results suggest that troriluzole reduces neuronal VGlut1 levels (**Fig. 2**). Comparing the IHC results with the cell culture results suggests that troriluzole may inhibit the pathway by which P301L tau mediates an increase in VGlut1 (Siano *et al*. 2023). It is now established that P301L blocks the function of the epigenetic factor HDAC1, which leads to an increase in VGlut1 expression per vesicle (Siano *et al*. 2023). This result provides new insight into the pathway by which troriluzole may prevent the increase in VGlut1. Because the number of VGLUT molecules has a direct impact on the number of glutamate molecules released by a single synaptic vesicle during exocytosis (Herzog *et al*. 2006; Wilson *et al*. 2005), altering VGlut levels will influence presynaptic release to ultimately change the levels of glutamate in the extracellular space. This is supported by our findings of restored presynaptic release probability (**Fig. 1G & 4B**), KCl-evoked release (**Fig. 3C**), and basal (tonic) glutamate levels (**Fig. 3A**) in 3xTg mice exhibiting increased VGlut1 levels that are reduced by troriluzole. These findings suggest that reducing VGlut1 expression can reduce extracellular glutamate levels. However, riluzole has also been demonstrated to improve glutamate uptake (Deng *et al*. 2009; Fumagalli *et al*. 2008). Though we did not observe differences in glutamate transporter 1 (GLT-1) expression (**Supp. Figure 1**), we cannot rule out that the reduction in basal glutamate observed in troriluzole-treated mice may be partially due to increased glutamate uptake.

In addition to restoring VGlut1 and glutamate levels, troriluzole treatment also restored the basal synaptic transmission (**Fig. 4A**) and synaptic plasticity deficits (**Fig. 4D**) observed in 3xTg mice. In addition, the anxiety-like phenotype in the elevated plus maze (**Fig. 6D**) observed in 3xTg, as well as the deficits in spatial reference memory in the Morris water maze, were rescued by treatment with troriluzole (**Fig. 7D**). Interestingly, we did not observe differences in the MWM during hidden platform testing, despite robust differences during probe trials assessing spatial reference memory. This suggests that the acquisition of the hidden platform was made using a non-spatial strategy in 3xTg mice. This is in line with prior studies showing that 3xTg mice tend towards non-spatial search strategies, including a chaining search strategy that is effective when a hidden platform is available (Baeta-Corral & Giménez-Llort 2015; Castillo-Mariqueo & Giménez-Llort 2020). In addition, 3xTg mice have exhibited hyperactive patterns in the MWM that have interfered with probe performance (Baeta-Corral & Giménez-Llort 2015), which may be related to the increased swim speed of 3xTg observed in the current study, which was also rescued by troriluzole treatment (**Fig. 7A**).

## Conclusion

This is the first study examining the effects of troriluzole in a preclinical AD model, and our findings support the use of troriluzole as an early therapeutic intervention. As a rationally designed prodrug of riluzole, troriluzole was engineered to overcome some of the pharmaceutical limitations of riluzole, including its low bioavailability, variable pharmacokinetic profile, negative food effect, requirement for twice daily dosing, and potential for hepatotoxicity (Pelletier *et al*. 2018; Pelletier *et al*. 2014; McDonnell *et al*. 2012). Overall, these results support the promise of glutamate-modulating drugs, such as troriluzole, in Alzheimer’s disease.

## Author contributions

J.P., P.D.P., S.P., E.R., L.C., M.W.G., and M.N.R. performed the experiments and/or analyzed the data. V.S., I.A.Q., M.W.G., and M.N.R. conceived and designed the experiments. All authors provided input in data interpretation and manuscript review. All authors have read and agreed to the published version of the manuscript.

## Data availability statement

The data that support the findings of this study are available from the corresponding author upon reasonable request.

## List of abbreviations

AD: Alzheimer’s disease
Aβ: β-amyloid
SVs: synaptic vesicle
MCI: mild cognitive impairment
APPswe: amyloid precursor protein Swedish
MAPT: microtubule-associated protein tau
TgNeg: Nontransgenic mice
DIV: days in vitro
Tro: troriluzole
Veh: vehicle
MWM: Morris water maze
GluOx: glutamate oxidase
DG: dentate gyrus
CA3: cornu ammonis 3
CA1: cornu ammonis 1
MEAs: microelectrodes arrays
fEPSP: field excitatory postsynaptic potentials
LTP: long-term potentiation
PPF: paired pulse facilitation
RMANOVA: repeated measures ANOVA
KS: two-sided Kolmogorov-Smirnov

