## Supplementary figures and images for "Troriluzole Rescues Glutamatergic Deficits, Amyloid and Tau Pathology, and Synaptic and Memory Impairments in 3xTg-AD Mice"

### Supplementary Figure 1

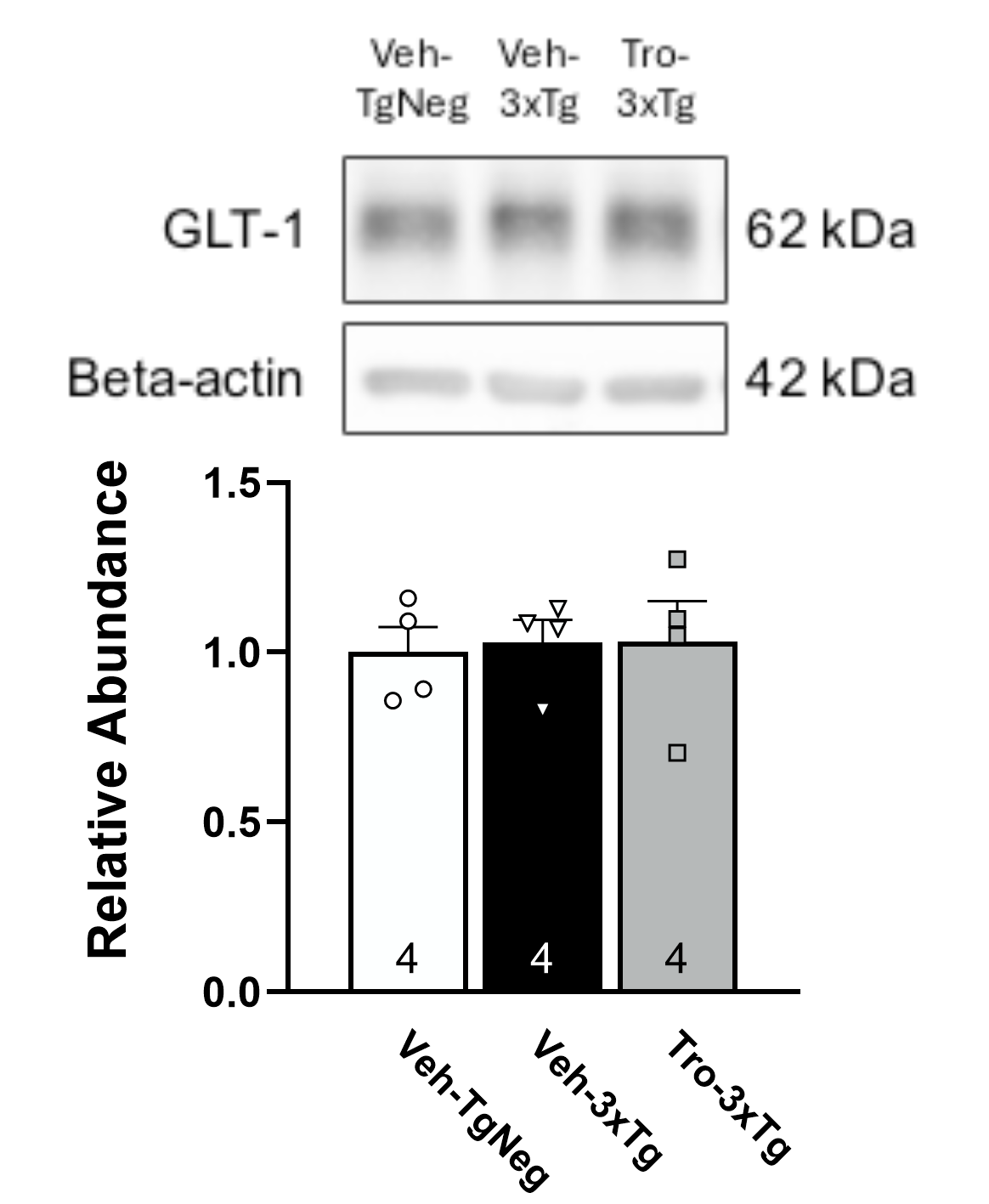
